# Somatic nuclear mitochondrial DNA insertions are prevalent in the human brain and accumulate over time in fibroblasts

**DOI:** 10.1101/2023.02.03.527065

**Authors:** Weichen Zhou, Kalpita R. Karan, Wenjin Gu, Hans-Ulrich Klein, Gabriel Sturm, Philip L. De Jager, David A. Bennett, Michio Hirano, Martin Picard, Ryan E Mills

**Affiliations:** Department of Computational Medicine and Bioinformatics, University of Michigan Medical School, Ann Arbor, MI 48109, USA; Department of Psychiatry, Division of Behavioral Medicine, Columbia University Irving Medical Center, New York, USA; Center for Translational & Computational Neuroimmunology, Department of Neurology, Columbia University Irving Medical Center, New York, NY 10032 USA; Taub Institute for Research on Alzheimer’s Disease and the Aging Brain, Columbia University Irving Medical Center, New York, NY 10032 USA; Department of Biochemistry and Biophysics, University of California, San Francisco, San Francisco, CA 94158, USA; Rush Alzheimer’s Disease Center, Rush University Medical Center, Chicago, IL 60612 USA; Department of Neurology, H. Houston Merritt Center, Columbia University Translational Neuroscience Initiative, Columbia University Irving Medical Center, New York, USA; New York State Psychiatric Institute, New York, USA; Department of Human Genetics, University of Michigan Medical School, Ann Arbor, MI 48109, USA

## Abstract

The transfer of mitochondrial DNA into the nuclear genomes of eukaryotes (Numts) has been linked to lifespan in non-human species ^1–3^ and recently demonstrated to occur in rare instances from one human generation to the next ^4^. Here we investigated numtogenesis dynamics in humans in two ways. First, we quantified Numts in 1,187 post-mortem brain and blood samples from different individuals. Compared to circulating immune cells (n=389), post-mitotic brain tissue (n=798) contained more Numts, consistent with their potential somatic accumulation. Within brain samples we observed a 5.5-fold enrichment of somatic Numt insertions in the dorsolateral prefrontal cortex compared to cerebellum samples, suggesting that brain Numts arose spontaneously during development or across the lifespan. Moreover, more brain Numts was linked to earlier mortality. The brains of individuals with no cognitive impairment who died at younger ages carried approximately 2 more Numts per decade of life lost than those who lived longer. Second, we tested the dynamic transfer of Numts using a repeated-measures WGS design in a human fibroblast model that recapitulates several molecular hallmarks of aging ^5^. These longitudinal experiments revealed a gradual accumulation of one Numt every ∼13 days. Numtogenesis was independent of large-scale genomic instability and unlikely driven cell clonality. Targeted pharmacological perturbations including chronic glucocorticoid signaling or impairing mitochondrial oxidative phosphorylation (OxPhos) only modestly increased the rate of numtogenesis, whereas patient-derived *SURF1*-mutant cells exhibiting mtDNA instability accumulated Numts 4.7- fold faster than healthy donors. Combined, our data document spontaneous numtogenesis in human cells and demonstrate an association between brain cortical somatic Numts and human lifespan. These findings open the possibility that mito-nuclear horizontal gene transfer among human post-mitotic tissues produce functionally-relevant human Numts over timescales shorter than previously assumed.

---

The incorporation of mitochondrial DNA into the nuclear genomes of organisms is an ongoing phenomenon ^4, 6–8^. These nuclear mitochondrial insertions, referred to as ‘Numts’, have been observed in the germline of both human ^6, 8–11^ and non-human ^7, 12–17^ species and occur as part of a wider biological process termed numtogenesis ^18, 19^, which has been defined as the occurrence of any mitochondrial DNA (mtDNA) components into the nucleus or nuclear genome. Once integrated, Numts are bi-parentally transmitted to future generations, like other types of genetic variation. While mostly benign, Numts have been implicated with cellular evolution and function ^1, 20^, various cancers ^18, 19^, and can confound studies of mitochondrial DNA heteroplasmy ^21, 22^, maternal inheritance of mitochondria ^10, 23–26^, and forensics ^27–29^.

Investigations of Numts have been conducted in numerous species, but yeast, in particular, has provided an excellent experimental platform as a model organism due to its smaller genome and fast replication timing. Mechanisms of Numt integration involve genome replication processes in several yeast species ^1, 2^ and have further been linked to the yeast *YME1* (yeast mitochondrial escape 1) gene and double-stranded break repair ^30–32^. Interestingly, Numts have also been associated with chronological aging in *Saccharomyces cerevisiae* ^3^, suggesting a model where the accumulation of somatic mutations with aging ^33, 34^, particularly structural genomic changes, could provide an opportunistic environment for somatic numtogenesis.

In humans, neural progenitor cells and cortical neurons harbor extensive tissue-specific somatic mutations, including single nucleotide variants (SNVs) ^35–37^, transposable elements ^38–40^, and larger structural variants ^41–43^. However, to date, no studies have investigated the extent of Numts specific in human brain regions, though several studies have now explored somatic numtogenesis in various cancers ^18, 44^. Using blood as the source of DNA, rare events of germline numtogenesis leading to a new Numt absent from either parents is estimated to occur every 4,000 human births, and to be more frequent in solid tumors but not hematological cancers ^4^. Using the observations in yeast as a foundation, we hypothesized that the accumulation of somatic mutations with age in the human brain also could be associated with numtogenesis and an increase in the number of somatically acquired (i.e., de novo) Numts. Mechanistically, numtogenesis requires the release of mitochondrial fragments into the cytoplasm and nucleus ^45, 46^, where they can be integrated into autosomal sequences. In this context, we note that neuroendocrine, energetic, and mitochondrial DNA maintenance stressors in human and mouse cells trigger mitochondrial DNA release into the cytoplasm ^47^ and even in the bloodstream ^48, 49^. Thus, intrinsic perturbations to mitochondrial biology or environmentally-induced stressors could increase numtogenesis across the lifespan.

We investigated these scenarios through a multifaceted approach using post-mortem human brain tissue and blood from large cohorts of older individuals, as well as a longitudinal analysis of primary human fibroblasts from healthy donors and patients deficient for *SURF1*, a gene associated with Leigh syndrome and cytochrome *c* oxidase deficiency ^50^ that alters oxidative phosphorylation (OxPhos). We further examined the potential role of environmental stress on numtogenesis through the treatment of these cells with oligomycin (OxPhos inhibitor) and dexamethasone (glucocorticoid receptor agonist).

## Results

### Somatic Numt integration differs by tissue, age, and cognitive status

We applied our Numt detection approach, *dinumt*, to WGS data generated in the ROSMAP cohort ^51, 52^ comprising 466 dorsolateral prefrontal cortex (DLPFC), 260 cerebella, 68 posterior cingulate cortex (PCC), and 4 anterior caudate (AC) tissue samples as well as non-brain tissue from 366 whole blood (WB) and 23 peripheral blood mononuclear cells (PBMCs) samples (**Methods**, **Fig. 1A**). We identified a mean of 10.4 Numts per sample across all tissues, of which ∼3 on average were found to be tissue-specific after filtering for germline Numt polymorphisms found in other samples, population-scale controls, or somatic Numts in other tissues. We observed no correlation between the number of Numts detected in each sample and its genomic sequence coverage (r^2^=0.003, **Supplementary Table 1**), indicating that our results are robust across a range of sequence depths ^53^. The detected Numts ranged from 20bp to 1,872bp in length, with a median of 63bp, a mean of 1,201bp, and s.d. 1,912bp (**Supplementary Table 1**), consistent with previous results from population-scale data in blood DNA ^4, 6, 7^.

**Figure 1.**
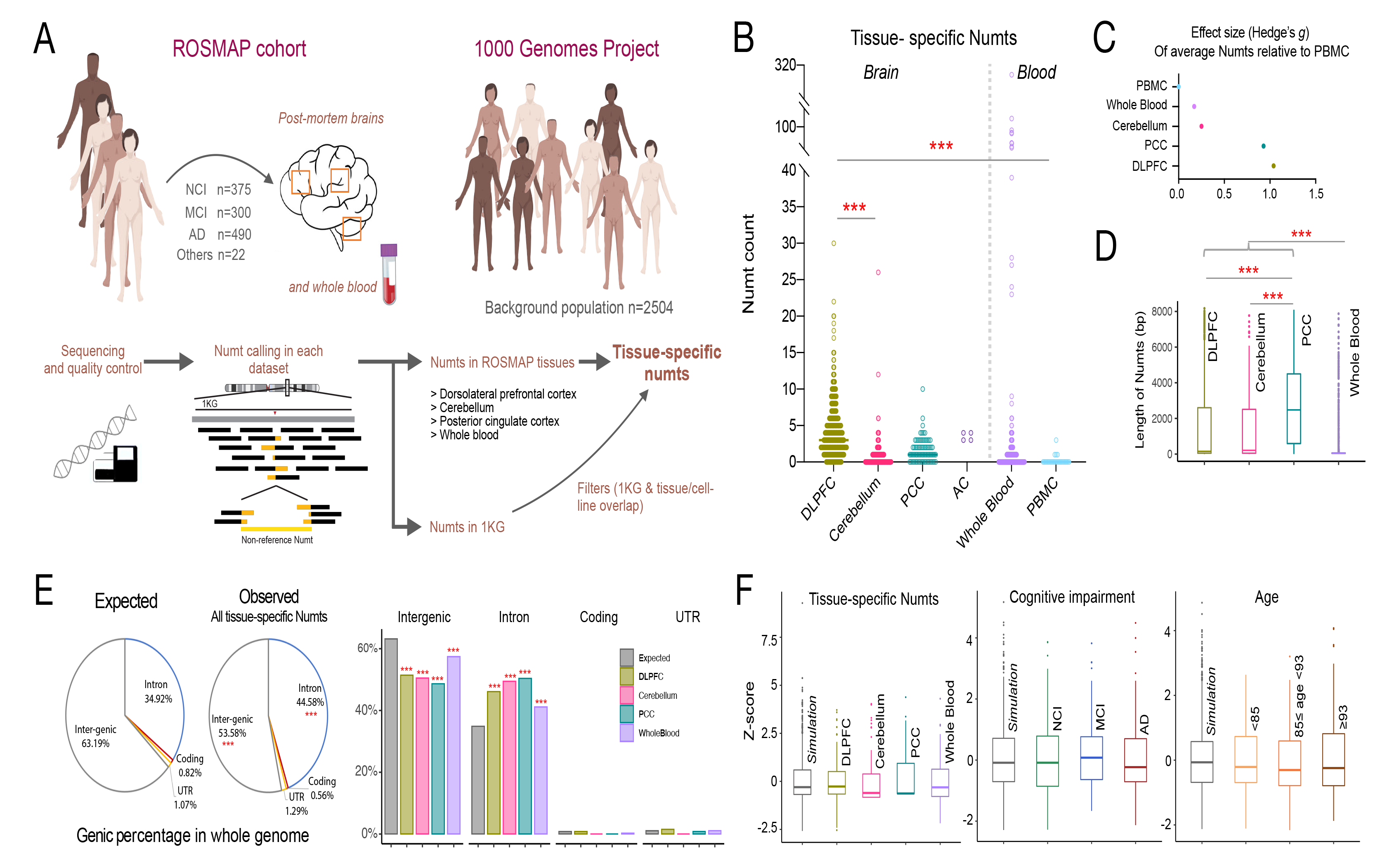
Characteristics of tissue-specific Numts in post-mortem human brain regions and blood. (A) Overview of approach to identify tissue-specific Numts in ROSMAP cohort. (B) Abundance of tissue-specific Numts across brain regions and blood cells from ROSMAP participants. (C) Effect size (Hedge’s *g*) of average tissue-specific Numts relative to peripheral blood mononuclear cells (PBMC). (D) Length of tissue-specific Numts across brain regions and whole-blood. (E) Genic distribution of tissue-specific Numts versus the expected distribution in the whole-genome for all samples (pie charts) and delineated by brain regions and whole-blood (bar charts). (F) Random genomic distributions of tissue-specific Numts across tissues (left), cognitive impairment stratifications (middle), and age groups (right), based on comparison to simulation data. The age groups were defined as less than 85 (n=295), between 85 to 93 (n=545), and greater than or equal to 93 (n=319). Student’s t-test was used to test the significance in B and D. Pearson’s chi-square test was used to test the significance in E. Fisher’s exact test was employed instead of Pearson’s chi-square test when sample sizes are small. ***, **, and * represent a significant p-value less than 0.001, 0.01, and 0.05, respectively.

We next examined whether there were differences in Numt abundance between tissues in aggregate, as few of the ROSMAP tissue samples were matched within the same individuals. We found the majority of tissue-specific Numts fell within DLPFC regions (mean=4.13 per person), representing a 5.5-fold (p-value<0.001) higher frequency compared to the cerebellum (mean=0.75), 2.4-fold higher than PCC (mean=1.71, p-value<0.001), and 15.9-fold higher than PBMC (mean=0.26, p-value<0.001) (**Fig. 1B**). Relative to PBMCs, DLPFC also showed the highest effect size (Hedge’s g=1.05) in Numt abundance, followed by PCC (g=0.94) (**Fig. 1C**). These results are comparable in tissue specificity with recent studies of human brain mosaicism for SNVs and other variant types ^35^. Interestingly, we found that the length of tissue-specific Numts in three brain regions (DLPFC, cerebellum, and PCC) were significantly longer than those observed in whole-blood (median=49bp, p-value<0.001), with PCC-specific Numts themselves exhibiting larger lengths (median=2,477bp) than DLPFC and cerebellum (median=152bp and 210bp, p-value<0.001, **Fig. 1D**). The absence of large Numts in blood immune cells could reflect negative selection against new Numts ^4^.

We next explored Numt integration using a gene-centric approach in the tissues with the largest number of samples (DLFPC, Cerebellum, PCC, and Whole Blood). We focused on the Numts that inserted in and around transcribed regions of the genome (introns, coding, UTR, and intergenic regions). Surprisingly, we found that our somatic Numts integrated into introns at a significantly higher rate compared to its overall composition rate of the genome (44.58% v.s. 34.92%, p-value<0.001), while a negative enrichment was observed in intergenic regions (53.58% v.s. 63.92%, p-value<0.001, **Methods**, **Fig. 1E**). Consistent significant differences in genic distribution were further observed across the various tissues (**Fig. 1E**), suggesting a moderate gene tolerance and potential negative selection of somatic Numt integration in the human brain and blood cells, consistent with recent studies ^4^.

We lastly hypothesized that the genomic distribution of somatic Numt integration sites may differ between tissues or across different age groups. We tested this hypothesis by randomly permuting the position of our observed Numts 50,000 times and assessing whether our observed integration sites differed significantly when compared within predefined 10Mb windows across the genome. After the multiple test correction (Benjamini-Hochberg Procedure), we observed no significant deviations from random for any of the tested tissues (**Fig. 1F**). We further stratified our results by age and cognitive status (**Methods**) and likewise observed no differences in genomic distribution. This is in agreement with previous studies that suggest Numt integration is a random occurrence ^6, 7, 11^, though we note that the paucity of Numts overall may be underpowered and preclude an accurate assessment at such broad regions across the genome.

### DLPFC-specific Numts are negatively associated with age at death in persons without cognitive impairment

On the basis of potential adverse genomic effects of Numts ^4, 45, 46^ and the results above, we hypothesized that tissue-specific Numts were associated with mortality and age at death, though this correlation might differ between tissues or clinical diagnoses. We first examined each tissue in aggregate and observed almost no correlation in Numt abundance with the age of death (**Fig. 2A, Supplementary Fig. 1**). Given the existence of mitochondrial DNA defects alterations in the human brain with cognitive decline ^54^, we next stratified individuals with tissue-specific Numts by their cognitive status into no cognitive impairment (NCI), mild cognitive impairment (MCI), and Alzheimer’s dementia (AD, COGDX score, **Methods**). We found that in DLPFC tissues, NCI individuals that carried more Numts died earlier (r^2^=0.094, p-value<0.001), with 2 additional Numt insertions observed per decade of life lost (**Fig. 2B**). MCI individuals exhibited a similar but lower negative association (r^2^=0.031, p-value<0.05). However, no correlation between Numts and age at death was observed in the AD group (r^2^=0.009, p-value=0.19). In the cerebellum, we observed similar patterns of correlation between Numts and age at death among cognitive groups, albeit with weaker correlations (NCI: r^2^=0.044, p-value<0.05; MCI: r^2^=0.007, p-value=0.514; and AD: r^2^=0.045, p-value<0.05, **Supplementary Fig. 2**). These results indicate that Numts are negatively associated with age at death in certain brain regions of non-AD individuals and suggest that the pathogenicity of AD is likely uncoupled from age-dependent Numt integration.

**Figure 2.**
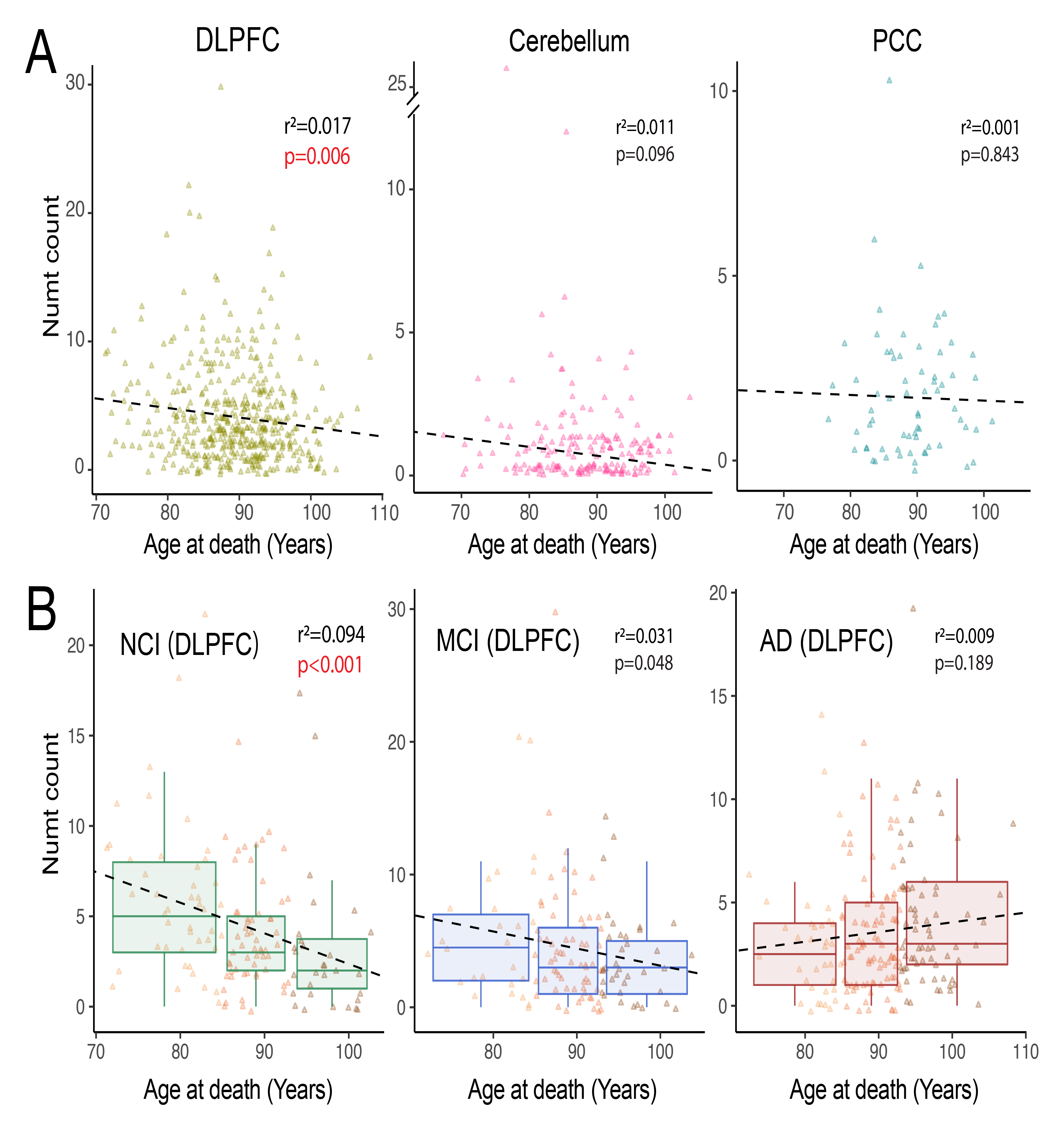
Numt association with age at death by tissue and cognitive impairment. (A) Correlation between age at death and abundance of DLPFC-specific, cerebellum-specific, and PCC-specific Numts, respectively. (B) DLPFC samples correlated with age at death, stratified by cognitive diagnosis status as no cognitive impairment (NCI, n=121, left), mild cognitive impairment (MCI, n=112, middle), and Alzheimer’s disease (AD, n=176, right). Data points are colored by arbitrary age groups (see **Methods**) in light yellow, orange, and brown, respectively. r^2^ and p-values are calculated using standard least-squares regression models.

### Somatic Numts accumulate over time in fibroblasts

The cross-tissue analysis of the ROSMAP cohort provided compelling results that Numts have both tissue and age-dependent characteristics of their integration in aggregate. However, the lack of relationships between individual tissue samples prohibits a direct measurement of Numt integration rates or numtogenesis. We therefore tested the dynamic transfer of Numts using a longitudinal, repeated-measures WGS study in primary human fibroblasts grown under physiological conditions (**Table 1**) ^5,55–57.^ Over time, replicating cells exhibit conserved epigenomic (hypomethylation), telomeric (shortening), transcriptional (senescence-associated markers), and secretory (pro-inflammatory) features of human aging, representing a useful model to quantify the rate of dynamic age-related molecular processes in a human system ^5^. We recently showed that primary mitochondrial bioenergetic defects accelerate the rate of aging based on the telomere shortening per cell division, DNA methylation clocks, and age-related secreted proteins^57^. Therefore, using this model to monitor the accumulation of overall and donor-specific unique Numts absent in the general population, we analyzed cultured fibroblasts from 3 unrelated healthy donors, aged in culture under physiological conditions for up to 211 days ^5^ (**Fig. 3A**). Instead of focusing on a single cell line tested in triplicates, we opted to include three separate donors, which provides a more robust test of our hypothesis.

**Figure 3.**
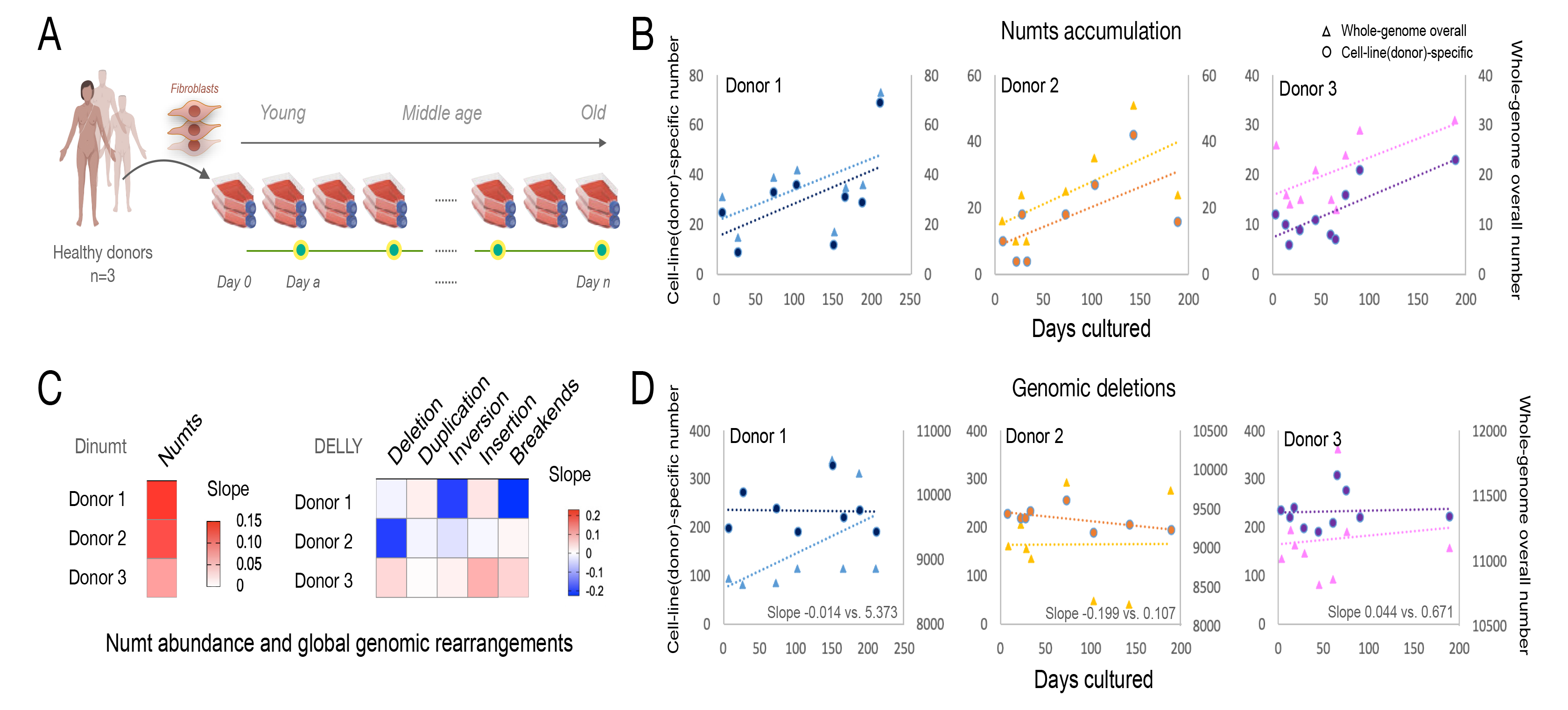
Numts accumulate in human fibroblasts during normal aging. (A) Study design of cellular aging model using primary fibroblasts from three healthy donors. (B) Numts accumulate over time in aging fibroblasts obtained from three healthy donors cultured up to 211 days, with left y-axis and circle dots representing cell-line specific variation, and right y-axis and triangle dots representing total accumulated variations from each cell line. (C) Heatmap of slopes based on the linear regression between days cultured and the cell-line specific calls, including Numts (from Dinumt, left) and structural variants (from DELLY, right). (D) Time-course of cell-line specific and whole-genome overall genomic deletions in the three healthy donors, with axis labels the same as in (B).

**Table 1.**
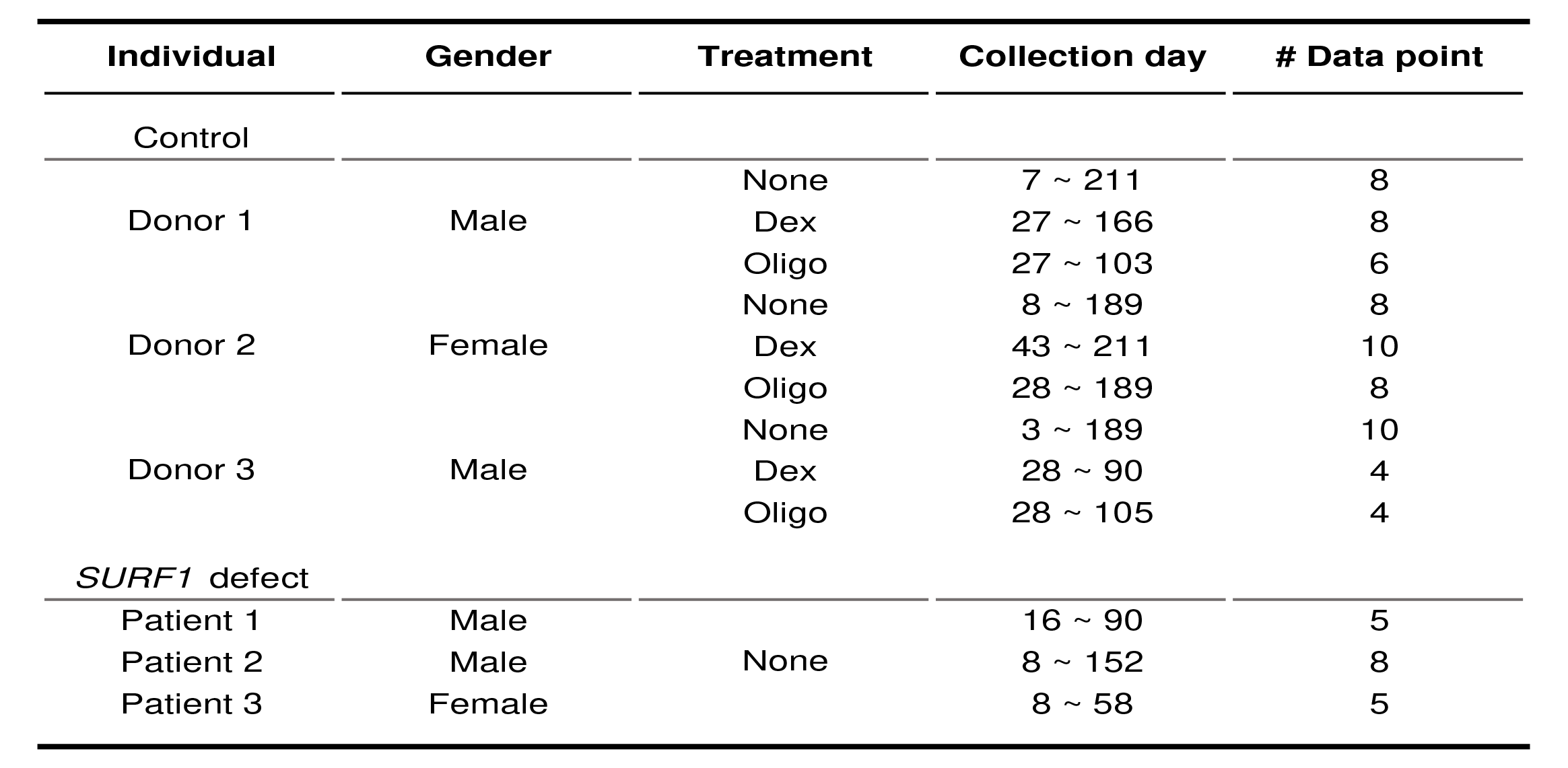
Experimental design for Numt accumulation study in the *in vitro* fibroblast aging model.

Across the three donors, we observed a positive correlation between time in culture and the number of unique Numts (r^2^=0.30, 0.38, and 0.59, respectively) with a positive slope range of 0.08-0.13 Numts/day. Thus, on average, human fibroblasts accumulated a novel Numt every 12.6 days of culture (or 0.79 Numt per 10 days, 95% C.I. = 0.28-1.31). Numtogenesis also was evident for both total Numts and cell-line specific Numt insertions (**Fig. 3B, Supplementary Fig. 3**). To determine if global genomic instability could account for this effect, we conducted the same analysis on multiple types of somatic structural variation (e.g., deletion, duplication, inversion, insertion, and breakends, **Methods**). The accumulation rates of these variant types was significantly lower compared to the rate of numtogenesis (**Fig. 3C**). For example, although we observed a positive yet more moderate increase in deletion abundance compared to Numts when considering all such variants, we did not observe the same increase in the cell-line specific deletions as with the somatic Numts (**Fig. 3D**). These results indicate that Numt insertions occur at a higher rate than autosomal deletions in this system and suggest a higher rate of age-related somatic numtogenesis rate than the other genetic variants (**Supplementary Fig. 4**). We further observed no significant increase in cell-line specific genomic duplications over time, thus indicating that the increase in the total number of Numts over time is likely due to novel integration events and not duplications of preexisting copies.

We questioned whether the accumulation of these apparently somatic Numts could be driven by the simple clonal expansion of few Numts-containing cells. Even within a given donor line followed longitudinally, all observed Numts were unique in their length and sequence (average 562bp and s.d. 1,400bp), and showed no evidence of relatedness with one another. This lack of sequence overlap is most parsimoniously explained by the random nature of our sequencing coverage (sequencing depth: 25x; total number of genomes in each experiment 2×10^6^ diploid genomes) and a large number of new Numts accumulating over time. Thus, the unique identity of all observed Numts in these *in vitro* experiments argues against the clonal origin of these events.

### Impact of environmental and genetic stress on somatic Numt integration rates

We next explored whether the cellular environment could impact somatic numtogenesis by testing if chronic exposure to a stress-mimetic or an inhibitor of mitochondrial OxPhos would alter the rate of Numt accumulation in otherwise healthy and aging fibroblasts. We analyzed human fibroblasts derived from the same three healthy donors described above that were treated with (a) the glucocorticoid receptor agonist dexamethasone (Dex, 100 nM) and (b) the ATP synthesis inhibitor oligomycin (Oligo, 100nM). Similar to the untreated donors (see **Methods**, r^2^=0.296, p-value=0.004 linear regression), both treatment groups exhibited an accumulation of new Numts over time (see **Methods**, linear regression for Dex group r^2^=0.588, p-value<0.001, for Oligo group r^2^=0.215, p-value=0.052). Compared to the untreated group (0.79 Numt per 10 days, 95% C.I. = 0.28- 1.31), Dex and Oligo treatments tended to increase the rate of numtogenesis to 1.07 Numt (95% C.I. = 0.65-1.49) and 2.15 Numt (95% C.I. = −0.02-3.27) per 10 days, respectively (**Fig. 4A, B, D**). Although these differences in effects did not reach statistical significance, the accumulation of Numts over time in these biologically independent experiments from those above further document Numtogenesis in aging human cells *in vitro*.

**Figure 4.**
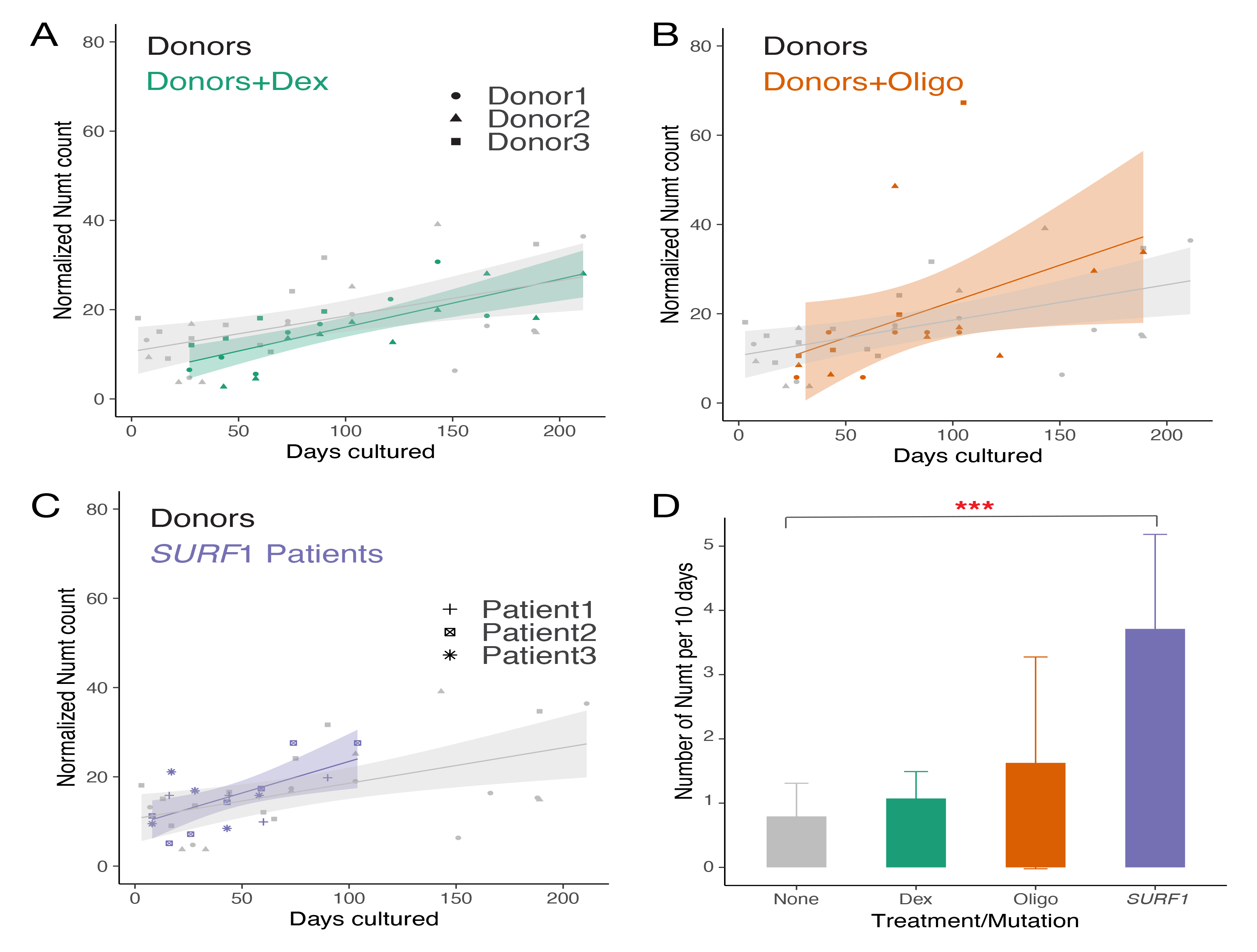
Effect of chronic pharmacological and genetic perturbation of mitochondrial function on Numt accumulation in fibroblasts. (A) Time course of Numt accumulation in healthy donors 1-3 and same cells cultured in dexamethasone (Dex) mimicking chronic Dex exposure. (B) Time course of Numt accumulation in healthy donors 1-3 and same cells cultured in oligomycin (Oligo). (C) Numt accumulation time course in the patient fibroblasts with *SURF1* gene defect (Patient 1-3) and the ones from three healthy donors (Donor 1-3). (D) Comparision of Slopes derived from all patients untreated, Dex-treated, oligo-treated, and *SURF1* gene defect. Numt counts for each group were normalized by the median value. A linear regression analysis was performed to derive the rate of the Numt accumulation and calculate the slopes in each group. ANOVA test was used to test the significance between the slopes of untreated donors and the ones of treated donors or patients in the hypothesis that pharmacological or genetic perturbation would increase the accumulation of Numts. ***, **, and * represent a significant p-value less than 0.001, 0.01, and 0.05, respectively, and ns represents non-significance.

Using a genetic approach, we further tested whether defects in mitochondrial OxPhos associated with mtDNA instability is sufficient to alter the rate of numtogenesis. We analyzed data from a similar fibroblast culture system in three patient-derived fibroblasts with *SURF1* mutations (Patient 1, 2, 3) ^5^. Mutations in *SURF1* represent one of the most frequent causes of cytochrome c oxidase and OxPhos deficiency in humans ^58^., and we recently showed that these *SURF1*-mutant fibroblasts accumulate large-scale mtDNA deletions over time in culture, demonstrating mtDNA instability in this model. In these independent donors, we again observed the accumulation of new Numts over time (regression r^2^=0.642, p-value<0.001). Strikingly, the rate of Numtogenesis in *SURF1*- mutant cells was 3.71 Numt per 10 days (95% C.I. = 2.24-5.18), in contrast to the rate of 0.79 Numt per 10 days in the healthy donors (4.7-fold of control, p-value=8.20E-05, ANOVA test, **Fig. 4C, D**). As in tumors ^18, 19^ and yeast ^1, 2^, this result further documents Numtogenesis as an inter-genomic event occurring in human cells over relatively short timescales, and establishes, using patient-derived cells with mtDNA instability, the modifiability of the rate of numtogenesis *in vitro* ^57^.

## Discussion

The transfer of mitochondrial DNA into the nuclear genome of eukaryotes occurs in the germlines of various eukaryotes, suggesting that the endosymbiotic event initiated 1.5 billion years ago is still ongoing ^4^. However, the extent and impact of somatic Numt insertions in specific human tissues have remained elusive outside of cancerous environments. Here, we provide some of the first evidence of the somatic nuclear colonization of Numts in both healthy and impaired human brain tissue across different age ranges. We found that specific brain regions harbor more somatic insertions than others, in line with previous studies of other types of genomic variation ^41–43^, and that these rates do differ with the degree of cognitive impairment. We further extend these observations in a longitudinal study of primary human fibroblasts under various environmental and genetic conditions, documenting variable rates of numtogenesis in dividing human cells.

Our data provide new information concerning the rate of numtogenesis in humans. While the cross-sectional study of post-mortem human brains does not allow us to draw conclusions about the rate of Numt transfer, the higher frequency of Numts in post-mitotic brain tissue - relative to the commonly studied genomic material from blood - indicates a greater number of Numts. On the other hand, our longitudinal in vitro studies allows us to measure Numts among the same cell population (i.e., “individual”) over time as cells accumulate age-related molecular hallmarks of aging. In this system, cells divide approximately every 40 hours (1.7 days) when they are young (from days 0-80) and slow their replication rate dramatically towards the end of life, when they undergo less than one replicative event per month (see growth curves in ^59^). Although the temporal resolution of our trajectories is limited by the number of timepoints across the lifespan of each cell line (on average 7), our data suggests that the rate of numtogenesis is roughly linear across the wide range of replication rate, and therefore more dependent on time rather than the rate of replication. This result is unlikely explained by clonality as all discovered Numts are unique. Each cell population contains between 1-5 million cells, meaning 2-10 million genomic copies. At 25x coverage, we therefore sample 0.00125- 0.00025% of all copies. Our sampling is therefore essentially random. If the accumulation of Numts in aging fibroblasts was driven by clonality we would see an increase in abundance of a few Numts present at each passage in the same donor, rather than an increased frequency of unique Numts at each time point, as observed here.

At least three main findings from our results and that of others suggest that de novo Numts may be functionally meaningful. *First*, strong evidence that *detectable* Numts are excluded from coding DNA sequences and instead preferentially integrate within intergenic regions ^4, 6–8^ suggest that they are under evolutionary constraint. In contrast, tumors frequently contain Numts within genes, and Numts may even contribute to oncogenesis, which in this case would drive positive selection in tumors ^4^. *Second*, Numts prevalence correlates with the somatic selection pressure for adverse mitochondrial genomic changes. In the replicating blood immune compartment of the bone marrow, high selection pressure occurs and eliminates mtDNA mutations over time ^60^, whereas de novo mtDNA defects accumulate at high levels in post-mitotic tissues such as skeletal muscle, heart and in the brain ^61^. Similarly, it is possible that Numt insertions which negatively affect cell fitness in the lymphoid or myeloid lineages of the bone marrow are outcompeted and eliminated from the cell pool (or exist at low levels and are not sampled during blood draw), compared to somatic tissues where the same potentially deleterious Numt insertions in post-mitotic cells cannot readily be outcompeted and eliminated without functionally compromising the tissue. *Third*, and relatedly, larger Numts that due to their size are more likely to be deleterious are observed across multiple post-mitotic brain regions compared to blood immune cells, which contain Numts approximately 1-2 orders of magnitude smaller than in brain samples.

Some limitations of our study should be noted. While our human multi-tissue study includes >1,000 individuals, given the low number of Numts per person, we are likely underpowered to draw definitive conclusions around the cartography of Numts, such as potential Numts hotspots in the nuclear genome. The report by Wei et al. in 66,083 human genomes robustly addresses this point ^4^. The absence of WGS data from multiple tissues in the same individuals also precludes direct comparisons of the specific rate of numtogenesis between tissues. This question could be addressed in other studies (e.g., GTEx) by sequencing dozens of tissues from the same individuals, but the current lack of such dataset precludes a robust analysis of this kind. In relation to our longitudinal cellular studies with genetic and pharmacological mitochondrial OxPhos defects, the marginal increase in cell death when OxPhos is disrupted ^57^ or with chronic glucocorticoid stress ^62^ could offer a route of negative selection that eliminates deleterious de novo Numts; the most compromised cells die, cleansing the cell population of new variants. If this effect was strong, it would make numtogenesis undetectable to WGS, or reduce the observed effect size of the rate of numtogenesis in all of our studies. This possibility could be addressed in future studies by systematically sequencing cellular debris or dead cells from the culture medium. Thus, while our positive results conclusively establish the dynamic nature of Numts transfer in healthy and stressed human cells, the magnitude reported across donors (range of 0.79 (controls) to 3.71 (*SURF1*-mutant) Numts every 10 days) may reflect a lower bound, and the rates of numtogenesis in cells under stress should be interpreted with caution. Validating the dynamic nature of numtogenesis across the lifespan in humans would require repeated measures, longitudinal WGS of post-mitotic tissues (e.g., muscle biopsies), with the caveat that the same exact cells likely cannot be repeatedly biopsied, and therefore that new Numts would be missed. For the reasons mentioned above (negative selection in the immune compartment), repeated WGS in blood would likely underestimate the true rate of *in vivo* mito-nuclear genomic transfer.

In conclusion, our results demonstrating the high prevalence of non-germline Numts in hundreds of human brains and their negative association with age at death suggests that numtogenesis occurs across the human lifespan and that they may have deleterious health effects. Using a longitudinal *in vitro* human system, we establish that primary fibroblasts accumulate Numts over time and that numtogenesis may be accelerated by some stressors, in particular *SURF1* defects associated with mtDNA instability. These findings build and extend previous evidence that numtogenesis is active in the human germline and can have deleterious genomic, cellular, and health effects on the host organism. The active transfer of mtDNA sequences to the nuclear genome adds to the vast repertoire of mechanisms of mito-nuclear communication that shape human health^63^.

## Methods

### ROSMAP cohort

#### Study participants

The Rush Memory and Aging Project (MAP) and the Religious Orders Study (ROS) ^51, 52^ are two ongoing cohort studies of older persons, collectively referred to as ROSMAP. The ROS study enrolls older Catholic nuns, priests, and brothers, from more than 40 groups across the United States. The MAP study enrolls participants primarily from retirement communities throughout northeastern Illinois. Participants in both cohorts were without known dementia at study enrolment and agreed to annual evaluations and brain donation on death. Studies were approved by an Institutional Review Board of Rush University Medical Center and all participants signed informed consent, Anatomical Gift Act, and a repository consent to share data and biospecimens.

The clinical diagnosis of AD proximate to death was based on the review of the annual clinical diagnosis of dementia and its causes by the study neurologist blinded for post-mortem data. Post-mortem Alzheimer’s disease pathology was assessed as described previously ^64, 65^ and Alzheimer’s disease classification was defined based on the National Institutes of Ageing-Reagan criteria ^66^. Dementia status was coded as no cognitive impairment (NCI), mild cognitive impairment (MCI), or Alzheimer’s dementia (AD) from the final clinical diagnosis of dementia and the NIA Reagan criteria as previously described ^67–69^.

#### Data processing

We obtained and processed whole genome sequence (WGS) samples from these cohorts through the NIA Genetics of Alzheimer’s Disease Data Storage Site (NIAGADS) data set NG00067. In brief, we obtained sequence data for 1187 tissue samples comprising 466 dorsolateral prefrontal cortex (DLPFC), 260 cerebella, 68 posterior cingulate cortex (PCC), 4 anterior caudate (AC), 366 whole blood (WB) samples, and 23 peripheral blood mononuclear cells (PBMCs). These were provided in CRAM format and aligned to the human genome reference (GRCh37) with an average read depth of 45X. Based on the clinical information of age, we also stratified all 1187 samples into three age groups with a similar sample size: a) samples died at an age younger than 85, b) age at death older than or equal to 85 and less than 93, c) age at death older than or equal to 93. All the alignment statistics are presented in **Supplementary Table 1**, along with the clinical characteristics of the study participants.

### In vitro fibroblast aging model

#### Fibroblast collection and passaging

We further made use of processed WGS generated from a recent study of aged primary human dermal fibroblasts ^5^. In brief, primary human dermal fibroblasts were obtained from distributors or our local clinic from 3 healthy and 3 *SURF1*-patient donors. Fibroblasts were isolated from biopsy tissue using standard procedures. Cells were passaged approximately every 5 days (+/- 1 day). Study measurements and treatment began after 15-day culture to allow for adjustment to the in *vitro* environment. Treatment conditions for healthy controls include the chronic addition of 1nM oligomycin (oligo) to inhibit the OxPhos FoF1 ATP synthase, and 100nM dexamethasone (dex) to stimulate the glucocorticoid receptor as a model of chronic stress ^62, 70^. Time points collected vary by assay, with an average sampling frequency of 15 days and 4-10 time points for each cell line and condition. Individual cell lines were terminated after exhibiting less than one population doubling over a 30-day period, as described in ^5^.

#### Whole-genome sequencing and processing

Whole-genome sequencing data were performed in the lifespan samples at each time point (overall 85 time points). Paired-end reads were aligned to the human genome (GRCh37) using Isaac (Isaac-04.17.06.15) ^71^. Samtools (Ver1.2) ^72^ and Picard Toolkit (http://broadinstitute.github.io/picard/) was further used to process the aligned bam files and mark duplicates. The average read depth from the WGS and other alignment statistics used in this study can be found in **Supplementary Table 2**.

### Population-scale WGS control data

We leveraged 2504 independent individuals from the 1000 Genomes Project Phase 3 to serve as population-level controls. Samples were sequenced by 30X Illumina NovaSeq (ftp://ftp.1000genomes.ebi.ac.uk/vol1/ftp/data_collections/1000G_2504_high_coverage/) ^73^, and the data were archived in CRAM format with the GRCh38 reference (ftp://ftp-trace.ncbi.nih.gov/1000genomes/ftp/technical/reference/GRCh38_reference_genome/). Numt calls in these samples were later converted to GRCh37 reference coordinates with the *liftOver* tool ^74^ for comparison with ROSMAP and lifespan data.

### Detection of non-reference Numts

We applied an updated version of Dinumt ^6, 7, 75^ to identify non-reference Numts across the different sequenced cohorts. Dinumt is an established software that was first used in the 1000 Genomes Project and validated by orthogonal methods, including PCR and Sanger sequencing ^6^, and long-read sequencing ^7^. Briefly, Dinumt identifies aberrant/discordant reads aligning with either the mtDNA or the reference Numts on one end and map elsewhere in the genome on the other end, read orientation, and various other filters to define insertion breakpoints. Reads are discarded if they don’t align uniquely to the nuclear genome and have a mapping quality (MAPQ) of less than 10. Identified insertions are then filtered for quality using a Phred scale (≥50), a cutoff of supporting reads (≥4), and a cutoff of read-depth (≥5X) around the insertion point. We built the first set of Numts as populational and polymorphic controls from the 2504 individuals of the 1KG Project recently re-sequenced to high coverage ^73^. Individual non-reference Numt callsets were resolved and merged into a single VCF file using the merging module of Dinumt. All ‘PASS’ Numts are lifted over ^74, 76^ from GRCh38 to GRCh37 for the downstream analysis. Dinumt was used to identify non-reference Numts across the individual sequences from 1187 samples in ROSMAP or 85 cell-line genomes in the lifespan model. The same criteria were conducted in the pipelines (**Fig. 1A**).

We identified tissue/cell-line specific Numt insertions using the identified non-reference Numts. Tissue-specific Numts were derived from the ROSMAP callset, and cell-line specific calls were derived from the Numt callset of fibroblast lifespan data. Numts from each sample were first merged into an aggregated set for these two callsets. We then extracted all non-reference Numts that were found in only one specific tissue across all samples or cell-line respectively (**Supplementary Table 1, 2**). All analysis pipelines and the command lines for running Dinumt can be found at https://github.com/mills-lab/numts- and-aging-in-fibroblasts-and-brains ^77^.

### Statistical analysis

Cell-line specific Numts from fibroblast lifespan data were grouped by both donor/patient and treatment status. There are four treatment statuses: no treatment donors, donors/cells cultured in dexamethasone (Dex), donors/cells cultured in oligomycin (Oligo), and patients’ fibroblasts with *SURF1* gene mutation. To increase the statistical power of data points in each group, we normalized the Numt count, merged the data points from individual samples, and then conducted the linear regression. Values of Numt Numbers were normalized by the median of Numt count in each category. A Linear regression model was constructed for each category respectively as below:

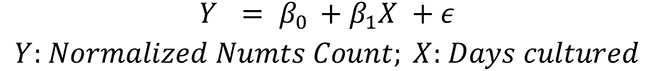

Slopes (*β*_1_) were compared by ANOVA test separately between two categories. All statistical analyses were performed in R 4.0.5.

### Genomic analyses for non-reference Numts

#### Numt hotspots across chromosomes

The entire nuclear genome was delineated into 10 Mbp bins and the frequency for tissue-specific Numts from ROSMAP in each bin was calculated. We performed a permutation analysis (with 50,000 simulation times matched to the number of Numt insertions) to determine any hotspots across genome bins compared to the real data. An empirical p-value was calculated for all the bins based on the frequency of Numt ranking in simulation data. A multiple test correction (Benjamini-Hochberg) was further conducted to decrease the false discovery rate. A bin with a p-value less than 0.05 after the adjustment was defined as a significant hotspot. We stratified the tissue-specific Numts into different tissues, cognitive impairment levels, and age groups to perform the hotspot analysis separately.

#### Genomic content analysis and functional annotation

We conducted the genomic content analysis for non-reference Numt insertions. We calculated the genic distribution for the tissue-specific Numts from ROSMAP. Gene track (GRCh37) was obtained from Ensembl Genome Browser (https://grch37.ensembl.org/). Parameters for protein-coding regions, transcriptomes, and exons were calculated based on a previous report ^78^. Pearson’s chi-square test and Fisher’s exact test were used to test the significant difference between the Numt genic distribution and reference genic distribution. GC content and repeat sequence analyses were carried out both in the set of cell-line specific Numt insertions from the lifespan model and polymorphic Numts from the 1000 Genomes project. GC content and repeat sequence were downloaded from the GC content table and RepeatMasker track in UCSC Genome Browser (https://genome.ucsc.edu/). Gene mapping was carried out by AnnotSV (https://lbgi.fr/AnnotSV/) ^79^ to determine the genes that were potentially affected by the tissue-specific Numts from ROSMAP (**Supplementary Table 3**) or cell-line specific Numts from the lifespan model (**Supplementary Table 4**).

### Detection of structural variation (SV)

Background structural variations (SVs) were detected in the data of the 1000 Genomes Project, ROSMAP, and lifespan model. We used an integrated non-reference SV callset from the 1000 Genomes Project as the control in the project to filter out potential non-somatic SVs at the population level. It was derived from 13 callers and can be obtained from http://ftp.1000genomes.ebi.ac.uk/vol1/ftp/data_collections/1000G_2504_high_coverage_SV/working/20210104_JAX_Integration_13callers/. Delly2 (Version 0.8.5) ^80^ was applied to resolve non-reference SVs (including deletions, duplications, insertions, inversions, and translocations), and MELT (Version 2.1.4) ^81^ was used to identify a specific type of non-reference SVs, mobile element insertions (MEIs, including Alus, LINE-1s, and SVAs), in the sequenced genomes of lifespan experiments. Manta ^82^ and Canvas (Version 1.28.0) ^83^ were also applied to resolve non-reference SVs in the sequenced genomes of lifespan experiments. The same pipeline used in Numts was implemented to identify tissue-specific or cell-line specific SVs/MEIs among the ROSMAP and lifespan samples (**Supplementary Table 1, 2**).

### Mitochondrial DNA copy number

#### Estimation of mtDNA copy number from WGS (ROSMAP and lifespan)

The median sequence coverages of the autosomal chromosomes *covnuc* and of the mitochondrial genome *covmt* were calculated using R/Bioconductor (packages GenomicAlignments and GenomicRanges). Ambiguous regions were excluded using the intra-contig ambiguity mask from the BSgenome package. The mtDNAcn was z-standardized within each brain region and DNA extraction kit and then logarithmized. The normalization facilitated the combined analysis of the two different kits used for the DLPFC and resulted in approximately normal mtDNAcn measures ^54^. In the lifespan study, mtDNA copies per cell were calculated as the ratio between the median coverage of the mitochondrial chromosome and twice the median coverage of the autosomes (mtDNA/2 X nDNA) from uniquely mapped paired-end reads with proper read orientation.

## Data availability

Whole genome sequence of ROSMAP can be obtained through the NIA Genetics of Alzheimer’s Disease Data Storage Site (NIAGADS) data set NG00067 ^51, 52^. Whole genome sequences of aged primary human dermal fibroblasts can be found in a recent study ^5^. Illumina-sequenced 2504 independent individuals from the 1000 Genomes Project Phase 3 can be found at: ftp.1000genomes.ebi.ac.uk/vol1/ftp/data_collections/1000G_2504_high_coverage/ ^73^. The Numt callsets by Dinumt from 1000 Genomes Project (GRCh38) and from ROSMAP and Lifespan dataset (GRCh37), the SV callsets by DELLY, Manta, and Canvas from Lifespan dataset (GRCh37), and the MEI callset by MELT from Lifespan dataset (GRCh37) can be found at GitHub page: github.com/mills-lab/numts-and-aging-in-fibroblasts-and-brains.

## Code availability

Dinumt: https://github.com/mills-lab/dinumt ^75^. The scripts and command lines in the project can be found at the GitHub page: github.com/mills-lab/numts-and-aging-in-fibroblasts-and-brains, and ZENODO: https://zenodo.org/record/6977577 ^77^.

## Supplementary Materials

**Supplementary Table 1. Meta table for ROSMAP sequencing data, including alignment statistics, clinical information, and variant numbers.**

**Supplementary Table 2. Meta table for lifespan sequencing data, including alignment statistics, experimental information, and variant numbers.**

**Supplementary Table 3. Gene annotation for tissue-specific Numts from ROSMAP. Supplementary Table 4. Gene annotation for cell-line specific Numts from lifespan model.**

**Supplementary Figure 1. Distribution of tissue-specific Numts by size and association with age at death and AD pathology.** (A) DLPFC-specific Numts by size and stratified with cognitive status. (B) Cerebellum-specific Numts by size and stratified with cognitive status. (C) PCC-specific Numts by size and stratified with cognitive status. Student’s t-test was used to test the significance. ***, **, and * represent a significant p-value less than 0.001, 0.01, and 0.05, respectively.

**Supplementary Figure 2. Cerebellum-, PCC-, and whole-blood-specific Numts are not associated with the age of death or cognitive status.** (A) Cerebellum samples correlated with age at death, stratified by cognitive diagnosis status. (B) PCC samples correlated with age at death, stratified by cognitive diagnosis status. (C) Whole-blood samples correlated with age at death, stratified by cognitive diagnosis status. Data points are colored by arbitrary age groups (see Methods) in light yellow, orange, and brown, respectively. r^2^ and p-values are calculated using standard least-squares regression models.

**Supplementary Figure 3. Common fibroblast Numts are not abundant across the lifespan and do not associate with age.** (A) Numts shared between cell lines (donors) are not significantly correlated with aging. (B) Slopes from cell-line specific Numts and shared Numts in the lifespan model.

**Supplementary Figure 4. Background somatic SVs and MEIs during aging in primary human fibroblasts.** (A) Heatmap of slopes based on the linear regression between days cultured and the cell-line specific Numts (from Dinumt). (B) Heatmap of slopes based on the linear regression between days cultured and the cell-line specific MEIs (from MELT). (C) Heatmap of slopes based on the linear regression between days cultured and the cell-line specific SVs (from DELLY).

## Author contributions

M.P., R.E.M., K.R.K., and W.Z. conceived the project. H.K., P.L.D., and D.B. facilitated access to and expertise for ROSMAP samples. M.H. established and cultured cell lines and G.B. conducted fibroblast experiments. R.E.M. and W.Z. developed and adapted the Dinumt pipeline. W.Z., K.R.K, and W.G. performed computational analyses. All authors guided the data analysis strategy. R.E.M., W.Z., M.P., and K.R.K. wrote the manuscript. All authors critically read, edited, and approved the final manuscript.

