## Supplementary figures and images for "Somatic nuclear mitochondrial DNA insertions are prevalent in the human brain and accumulate over time in fibroblasts"

### Supplementary.Figures.pdf

Supplementary.Figure1

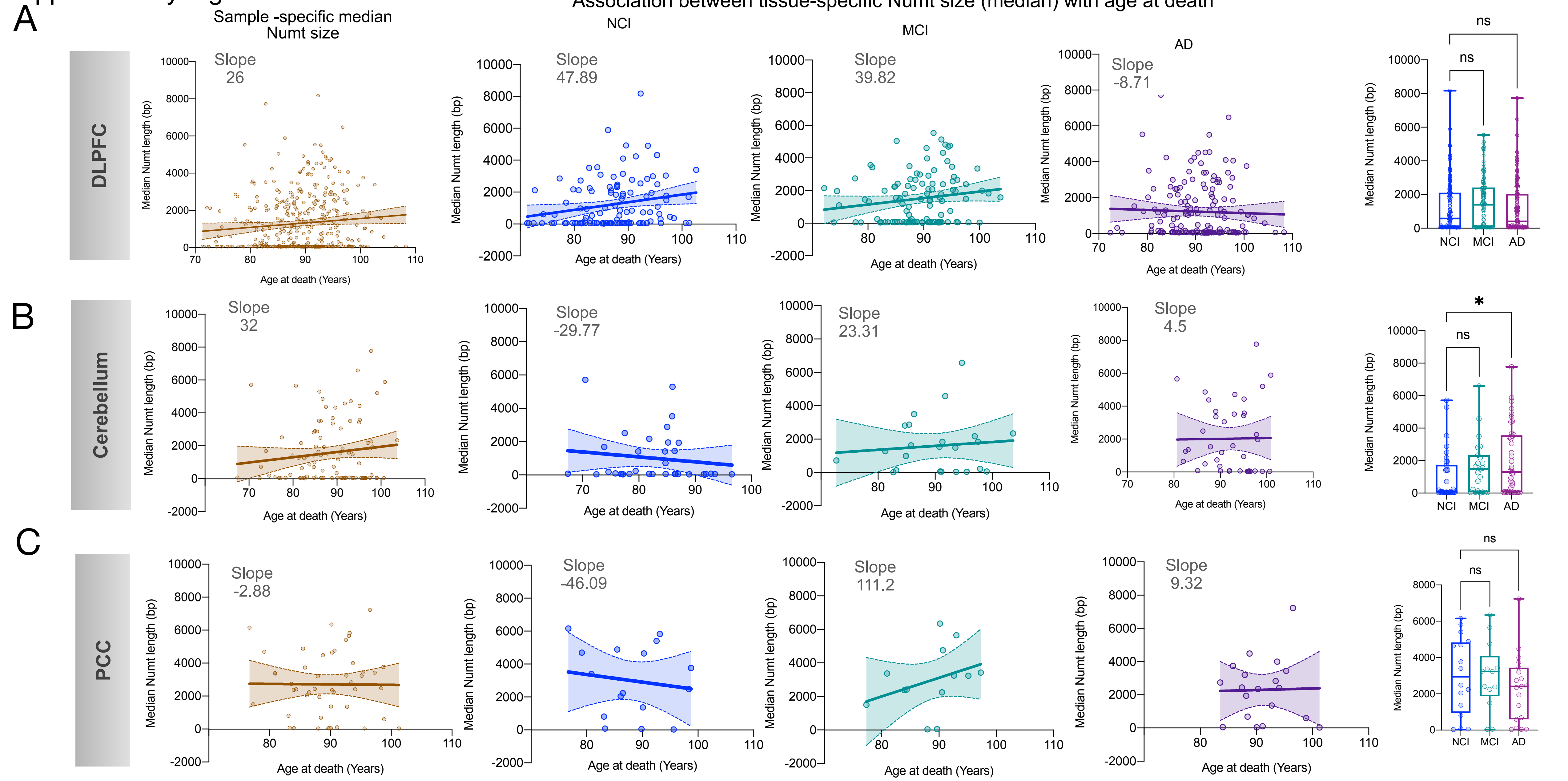

Supplementary.Figure2

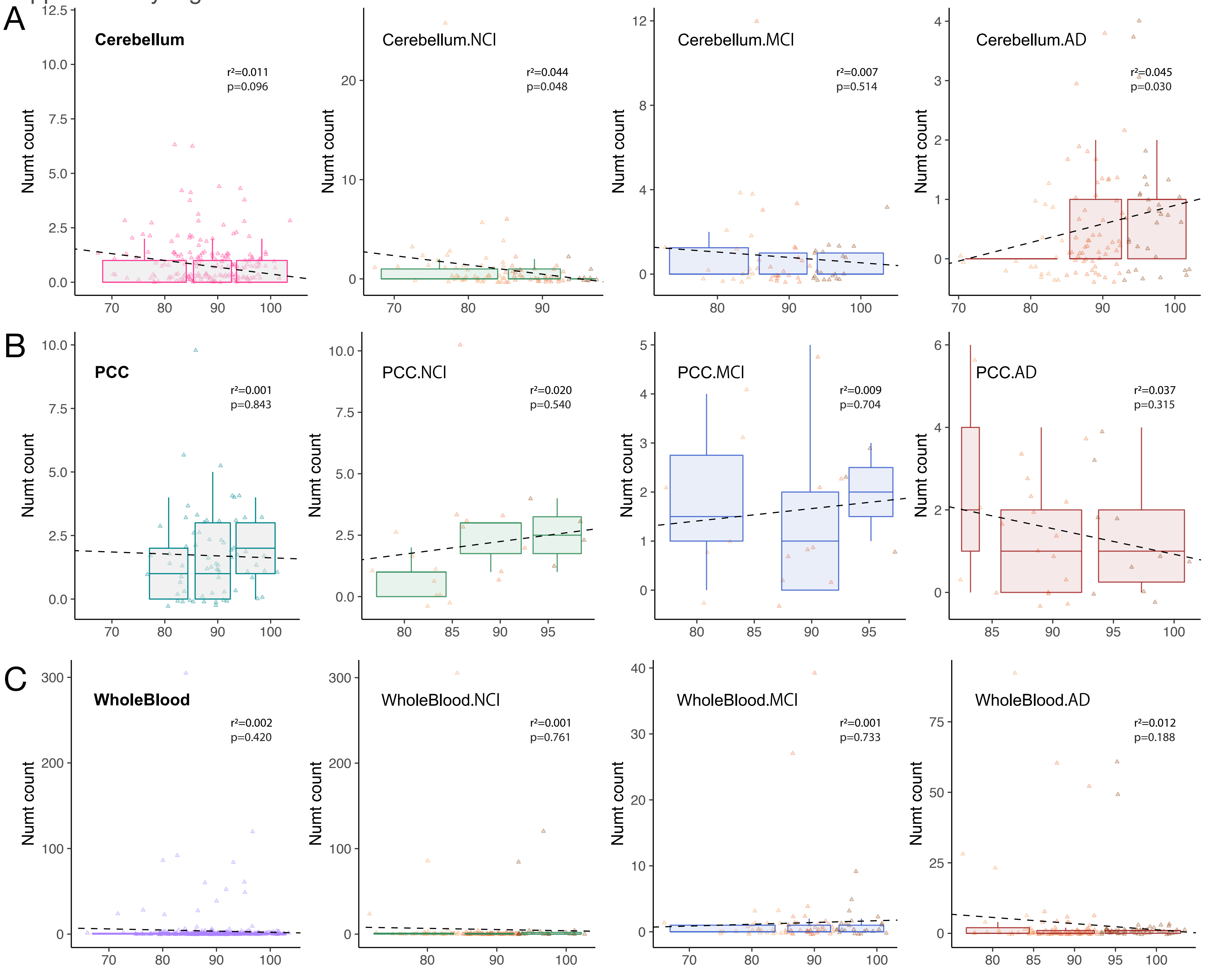

#### Supplementary.Figure3

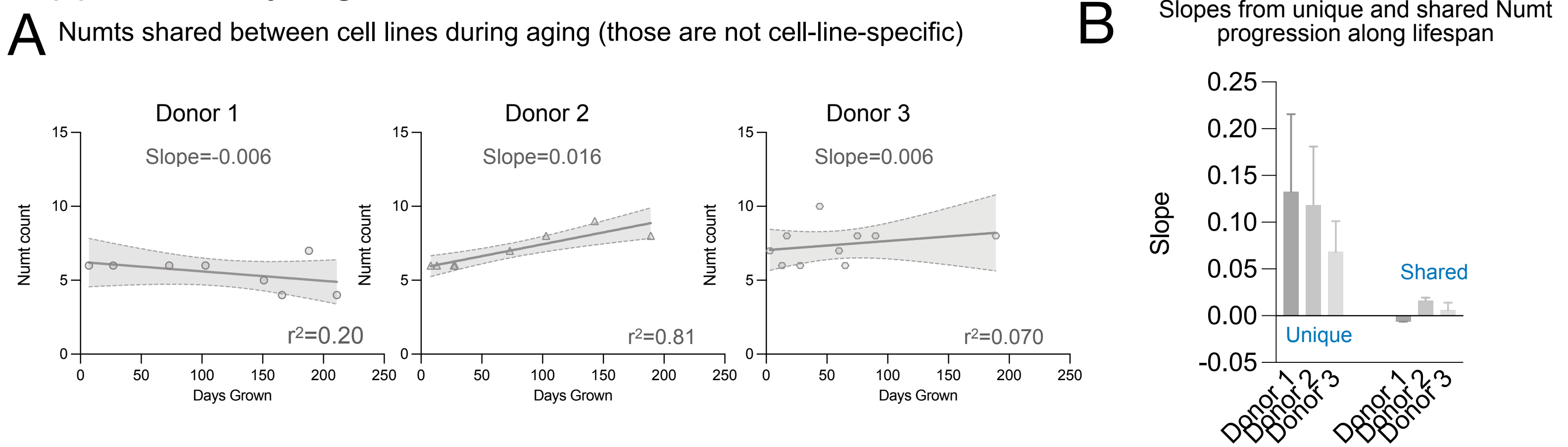

#### Supplementary.Figure4

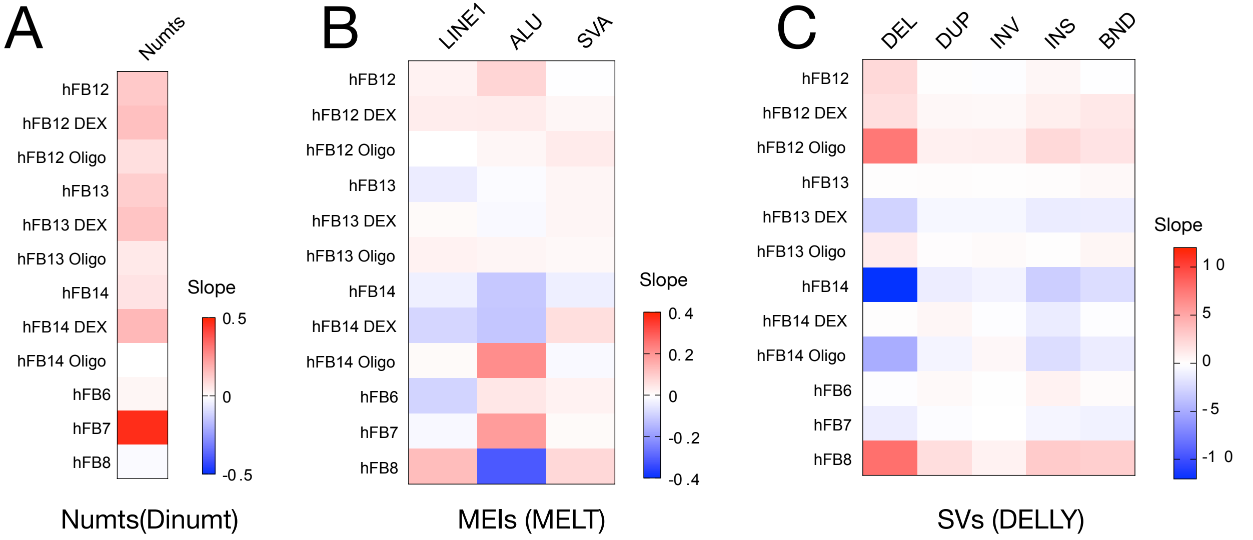
